# National and Regional Influenza-Like-Illness Forecasts for the USA

**DOI:** 10.1101/309021

**Authors:** Michal Ben-Nun, Pete Riley, James Turtle, David P. Bacon, Steven Riley

## Abstract

Health planners use forecasts of key metrics associated with influenza-like-illness (ILI); near-term weekly incidence, week of season onset, week of peak, and intensity of peak. Here, we describe our participation in a weekly prospective ILI forecasting challenge for the United States for the 2016-17 season and subsequent evaluation of our performance. We implemented a metapopulation model framework with 32 model variants. Variants differed from each other in their assumptions about: the force-of-infection (FOI); use of uninformative priors; the use of discounted historical data for not-yet-observed time points; and the treatment of regions as either independent or coupled. Individual model variants were chosen subjectively as the basis for our weekly forecasts; however, a subset of coupled models were only available part way through the season. Most frequently, during the 2016-17 season, we chose; FOI variants with both school vacations and humidity terms; uninformative priors; the inclusion of discounted historical data for not-yet-observed time points; and coupled regions (when available). Our near-term weekly forecasts substantially over-estimated incidence early in the season when coupled models were not available. However, our forecast accuracy improved in absolute terms and relative to other teams once coupled solutions were available. In retrospective analysis, we found that the 2016-17 season was not typical: on average, coupled models performed better when fit without historically augmented data. Also, we tested a simple ensemble model for the 2016-17 season and found that it underperformed our subjective choice for all forecast targets. In this study, we were able to improve accuracy during a prospective forecasting exercise by coupling dynamics between regions. Although reduction of forecast subjectivity should be a long-term goal, some degree of human intervention is likely to improve forecast accuracy in the medium-term in parallel with the systematic consideration of more sophisticated ensemble approaches.

**Author summary:** It is estimated that there are between 3 and 5 million worldwide annual seasonal cases of severe influenza illness, and between 290 000 and 650 000 respiratory deaths [1]. Influenza-like-illness (ILI) describes a set of symptoms and is a practical way for health-care workers to easily estimate likely influenza cases. The Centers for Disease Control (CDC) collects and disseminates ILI information, and has, for the last several years, run a forecasting challenge (the CDC Flu Challenge) for modelers to predict near-term weekly incidence, week of season onset, week of peak, and intensity of peak. We have developed a modeling framework that accounts for a range of mechanisms thought to be important for influenza transmission, such as climatic conditions, school vacations, and coupling between different regions. In this study we describe our forecast procedure for the 2016-17 season and highlight which features of our models resulted in better or worse forecasts. Most notably, we found that when the dynamics of different regions are coupled together, the forecast accuracy improves. We also found that the most accurate forecasts required some level of forecaster interaction, that is, the procedure could not be completely automated without a reduction in accuracy.

## Introduction

Infectious pathogens with short generation times pose public health challenges because they generate substantial near-term uncertainty in the risk of disease. This uncertainty is most acute and shared globally during the initial stages of emergence of novel human pathogens such as SARS [2], pandemic influenza [3], or the Zika virus [4]. However, at national and sub-national levels, uncertainty arises frequently for epidemic pathogens such as seasonal influenza, dengue, RSV and rotavirus; causing problems both for health planners and at-risk individuals who may consider changing their behavior to mitigate their risk during peak periods.

Seasonal influenza affects populations in all global regions and is forecast annually in temperate populations, either implicitly or explicitly [5]. Peak demand for both outpatient and inpatient care is driven by peak incidence of influenza in many years [6]. Therefore, the efficient provision of elective procedures and other non-seasonal health care can be improved by accurate forecasts of seasonal influenza. Implicitly, most temperate health systems use knowledge of historical scenarios with which to plan for their influenza season. The current situation is then assessed against the deviation from the historical averages and worst-cases as observed in their own surveillance system.

The United States Centers for Disease Control (CDC) has sought to formalize regional and national forecasts by introducing an annual competition [7]. Each week, participating teams submit weekly estimates of incidence for the next four weeks, season onset, and timing and intensity of the peak. Methods used by teams include purely statistical models, [8–10] mechanistic models [11,12] machine learning and hybrid approaches [13–15]. Expert-opinion surveys have also been used and performed well. Some teams augment their forecasts of the official ILI data with the use of potentially faster or less-noisy datasets such as google flu trends [16].

Here we describe a our mechanistic-model-supported participation in the 2016-17 CDC influenza forecasting challenge, as an example of a disease forecasting process. We emphasize a subjective human component of this process and also describe a retrospective evaluation of the models for the previous six seasons. All the models described are implemented in the R package Dynamics of Interacting Community Epidemics (DICE, https://github.com/predsci/DICE).

## Methods

### Data

The CDC Influenza-like-illness Surveillance Network (ILINet) Human and Health Services (HHS) region and national data were downloaded from the CDC-hosted web application FluView [17] and used to create a historic database of ILI cases. Fig S1 shows which states are grouped into each HHS region, along with the population of each region. Because we require an absolute number of cases per week, the CDC ILINet data is converted from percent ILI cases per patient to ILI cases. We estimate the absolute number of weekly ILI cases by dividing the weighted percent of ILI cases in the CDC data by 100 and multiplying it by the total weekly number of patients. We assume two outpatient visits per person per year so that the total weekly number of patients is estimated as: (total regional population)x(2 outpatient visits per person per year)x(1 year/52 weeks).

The estimate of two outpatient visits per year is based on two studies. In 2006 Schappert and Burt [18] studied the National Ambulatory Medical Care Surveys (NAMCS) and the National Hospital Ambulatory Medical Care Surveys (NHAMCS) and calculated an ambulatory rate of 3.8 visits per capita-year. The 2011 NAMCS [19] and NHAMCS surveys [20] estimated ambulatory visit rates of 3.32 visits per capita per year for physician’s offices, 0.43 for hospital emergency departments and 0.33 hospital Outpatient Departments. We sum these rates to get an outpatient visit per capita-year of 4.08. We further estimate from the surveys that only half of these outpatient clinics are sites that report to ILINet, and hence we rounded to our two outpatient visits per year estimate.

Specific humidity (SH) is measured in units of kg per kg and is defined as the ratio of water vapor mass to total moist air mass. Two other measurements of humidity are absolute humidity and relative humidity. SH is included in DICE as a potential modifier of transmissibility for this time period and uses Phase-2 of the North American Land Data Assimilation System (NLDAS-2) data base provided by NASA [21–23]. The NLDAS-2 data base provides hourly specific humidity (measured 2-meters above the ground) for the continental US at a spatial grid of 0.125^°^ which we average to daily and weekly values. The weekly data is then spatially-averaged for the states and CDC regions.

School vacation schedules were collected for the 2014-15 and 2015-16 academic years for every state. For each state, a school district was identified to represent each of the three largest cities. Vacation schedules were then collected directly from the district websites. These three school vacation schedules were first processed to a weekly schedule with a value of 0 indicating class was in session all five weekdays and a 1 indicating five vacation days. Next, the representative state schedule was produced by averaging the three weekly district schedules. Region schedules are obtained by a population-weighted average of the state schedules. Similarly, the national schedule is generated by a population-weighted average of the regions.

For the 2016-17 season we determine start and end times as well as spring and fall breaks from the previous years schedules. Thanksgiving and winter vacation timing was taken from the calendar where the winter break is assumed to be the last two calendar weeks of the year. Based on the proportion of schools closed and number of days closed, *p*(*t*) is assigned a value in the range [0, 1]. For example in week *t*_*i*_, if all schools are closed for the entire week then we define the proportion of open schools *p*(*t*_*i*_) = 1. However, if all schools have Monday and Tuesday off (missing 2 of 5 days), then *p*(*t*_*i*_) = 0.4. Similarly, if 3 of 10 schools have spring break (entire week off), but the other 7 schools have a full week of class then *p*(*t*_*i*_) = 0.3. If all schools have a full week of class then *p*(*t*_*i*_) = 0.

### Basic model

The DICE package has been designed to implement meta-population epidemic modeling on an arbitrary spatial scale with or without coupling between the regions. Our model for coupling between spatial regions follows ref [24]. We assume a system of coupled S-I-R equations (susceptible-infectious-recovered) for each spatial region. In this scenario, the rate at which a susceptible person in region *j* becomes infectious (that is transitions to the *I* compartment in region *j*) depends on: (1) the risk of infection from those in the same region *j*, (2) the risk of infection from infected people from region *i* who traveled to region *j*, and (3) the risk of infection encountered when traveling from region *j* to region *i*. To account for the three mechanisms of transmission, ref [24] defined the force of infection, or the average rate that susceptible individuals in region *i* become infected per time step as:

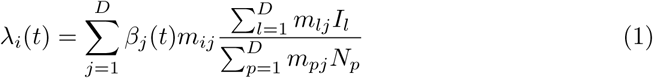

where *D* is the total number of regions. In our case, unlike reference [24], the transmissibility is not the same for all regions and it is allowed to depend on time: *β*_*j*_(*t*).

Given this force of infection we can write the coupled S-I-R equations for each region as:

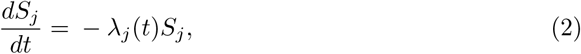

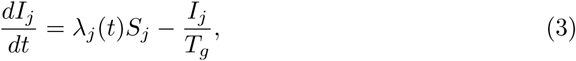

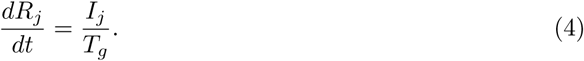

Eqs. (2–4) are the coupled version of the familiar S-I-R equations, where *T*_*g*_ is the recovery rate (assumed to be 2-3 days in the case of influenza). The mobility matrix, *m*_*ij*_, of Eq. 1 describes the mixing between regions. Thus, element *i, j* is the probability for an individual from region *i*, given that the individual made a contact, that that contact was with an individual from region *j*. As shown below, the sum over each row in the mobility matrix is one and in the limit of no mobility between regions the mobility matrix *m*_*ij*_ is the identity matrix so that 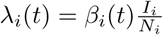 and we recover the familiar (uncoupled) S-I-R equations:

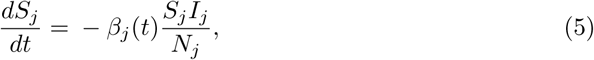

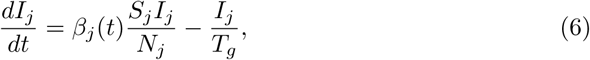

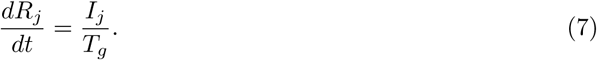

The level of interaction between spatial regions is determined by the mobility matrix and its interaction kernel, *κ*(*r*_*i*__*j*_):

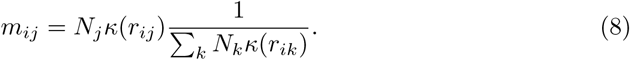

This kernel is expected to depend on the geographic distance between the regions (*r*_*ij*_), and following Mills and Riley [24] we use a variation of the off-set power function for it:

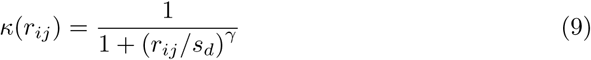

where *s*_*d*_ is a saturation distance in *km* and the power *γ* determines the amount of mixing between the regions: as *γ* decreases there is more mixing while as *γ* increases, mixing is reduced. In the limit that *γ → ∞* there is no mixing between regions and we recover the uncoupled SIR Eqs. (5–7). The DICE package is designed to allow the estimation of these two parameters (*γ* and *s*_*d*_), but they can also be set to fixed values.

The S-I-R equations model the total population, but the data are the number of weekly observed cases or incidence rate for each spatial region 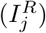. The weekly incidence rate is calculated from the continuous S-I-R model by discretizing the rate-of-infection term *λ*_*j*_(*t*)*S*_*j*_ (or 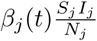 in the uncoupled case):

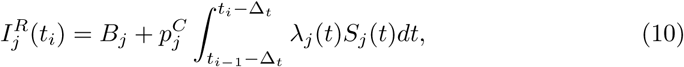

scaling by percent clinical 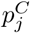, and adding a baseline *B*_*j*_. The term 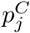 is the proportion of infectious individuals that present themselves to a clinic with ILI symptoms and *B*_*j*_ is a constant that estimates non-S-I-R or false-ILI cases. The integral runs over one week determining the number of model cases for week *t*_*i*_. Δ_*t*_ approximates the time delay from when an individual becomes infectious to when they visit a sentinel provider for ILI symptoms and is set to 0.5 weeks based on prior calibration [25,26]. Eq. 10 describes how DICE relates its internal, continuous S-I-R model to the discrete ILI data. In the next section we describe the procedure used for fitting this property (by optimizing the parameters: *β*_*j*_, *s*_*d*_, *γ*, *B*_*j*_, and 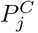) to an ILI profile.

To allow for different models for the force of infection/contact rate, we write this term in the most general way as a product of a basic force of infection, 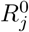, multiplied by three time dependent terms:

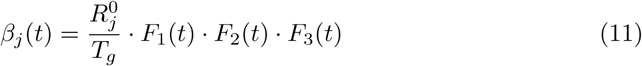

The first time dependent term, *F*_1_(*t*), allows for a dependence of the transmission rate on specific-humidity, the second (*F*_2_(*t*)) on the school vacation schedule, and the third (*F*_3_(*t*)) allows the user to model an arbitrary behavior modification that can drive the transmission rate up or down for a limited period of time. For the purpose of the CDC challenge we only considered models involving either *F*_1_(*t*), *F*_2_(*t*), both, or none (i.e., the contact rate does not depend on time), and the functional form of these terms is discussed in sections S1 Text and S2 Text of the Supporting Information.

### Fitting the model

The DICE fitting procedure determines the joint posterior distribution for the model parameters using a Metropolis-Hastings Markov Chain Monte Carlo (MCMC) procedure. [27] We describe the procedure starting with the simpler uncoupled case. In the uncoupled scenario infection can only happen within each HHS region, since there is no interaction between different spatial regions. The uncoupled regions are run sequentially and posterior distributions for the model parameters and forecasts are obtained. For each region, we simulated three MCMC chains each with 10^7^ steps and a burn time of 2 × 10^6^ steps. The smallest effective sample size that we report for any parameter is greater than 100. After sampling from the individual posterior densities of each region, we calculate our national forecast as the weighted sum of the regional profiles with the weights given by the relative populations of the regions. The national curve was also fitted directly (without any regional information) using all the models and priors, but these direct results were only used at the end of the season when estimating the performance of each of our procedures.

In the coupled scenario, the MCMC procedure uses Eqs. (2–4) along with Eq. (10) to simultaneously generate candidate profiles for the coupled ten HHS regions. The log-likelihood of the ten regional profiles is calculated and combined with the proper relative weights to generate a national log-likelihood which is minimized. It is important to note that in the coupled scenario we *only* optimize the national log-likelihood, and not the individual region-level likelihoods, but the parameters we optimize are still mostly region specific (only *s*_*d*_ and *γ* are not). We also tried fitting the coupled model to the regional log-likelihoods, however the results of the fits were not as accurate as the ones obtained when the national likelihood is optimized (see Discussion).

Both the coupled and uncoupled scenarios begin with the entire population of each region, minus the initial seed of infections which were fitted, in the susceptible state. We also fitted the onset time of each local epidemic and the proportion of infections that were cases. We also considered fitting an initially removed fraction *R*(0), but found that *p*_*c*_ and *R*(0) were very strongly correlated if both were fitted parameters.

### Specific model variants

We ran the model using four variants for the force of infection: (i) The force of infection depended on specific humidity only (H), (ii) school vacation only (V), (iii) both (HV) or, (iv) none (F). In all four variants we fit 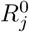 and fix *T g* at a value of 3 days. The allowed range for 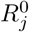 is between 1 and 3 with typical values being in 1.1 − 1.3.

We compare the performance of the model variants to: (1) Each other; (2) An ensemble model which is the equally weighted average of all the model variants; and (3) A historic NULL model, calculated as the weekly average of the past ten seasons (excluding the 2009 pandemic season).

#### Informative priors

In the previous section we described a traditional MCMC procedure which uses a log uniform distribution for the parameters, which we term an uninformative prior (UP). Early in the flu season, before the ILI curve takes off, this fitting can result in peak intensities that are significantly larger/lower than expected (based on historic values) and/or peak weeks that are inconsistent with past values. One way to constrain the predictions, which has been used by others [28,29], is to use an informed prior (IP).

To generate informative priors, we used each of the models supported by DICE to fit all previous seasons (starting from 2004) at both the national and regional levels. Using the history of the MCMC chain we then built a posterior distribution for each parameter and fitted it to a 1-D Gaussian. Although it would have been possible to use the posteriors directly from prior years, and thus preserve correlation structure in those distributions, we chose not to do this for two reasons. First, for the parameters we were informing, there was little correlation. Second, by fitting to a 1-D Gaussian, we were able to state precisely the prior assumptions for our actual forecast and quickly assess the degree to which they are informative. This would have been more difficult had we used stochastic draws from multivariate posteriors generated for previous years.

By repeating this procedure for each season and each model, we create a database of prior knowledge which can be used to inform the MCMC procedure for the current season. Specifically, at each week during the CDC challenge, and for each of the ten HHS regions and the national, we find the past season that is most similar to the current data (based on the value of the Pearson correlation, calculated using all weeks of available data and typically found to have a p-value that is > 0.9) and use the posterior distributions of that season as an informed prior for the current season. Each region has its own informed prior which is allowed to change from one epidemic week to the other. The Gaussian informed prior provides a simple way to penalize the likelihood when the parameter values sampled by the MCMC procedure are far from what was observed in a past similar season. To allow for an informed prior that is less restrictive, we also use a heated informed prior, where the Gaussian temperature is increased by an order of magnitude (which is equivalent to increasing the variance by a factor of ten). In the Results section we refer to the fitting procedures that use a prior as IP and HIP for informed prior and heated informed prior, respectively. Informed priors were used *only* with the uncoupled SIR Eqs. In a future study we plan to explore how they would extend to the coupled MCMC procedure.

#### Using discounted historical data for not-yet-seen future time points

In addition to informative priors, we also used data augmentation to make maximum use of prior data within a mechanistic framework. For each week during the challenge, our data augmentation was a form of extrapolation in which future unobserved time points were assumed to take either a historical average or values equal to those in the most similar prior season. However, these historically augmented time points were not counted within the likelihood with the same weight as actual observations. The weighting was equal to the value of the Pearson correlation between the observed data in the current year and the historical data for the same period from the year used for augmentation. We shifted from the historic data to the most similar data at epidemic week 6 (EW06) when we subjectively determined that the current season is very different from the historic average. The augmented data was also y-shifted so that it matched the last data point for the current season. The augmented data procedure was used for both the coupled and uncoupled fits and also using a heated augmented procedure (where the log-likelihood is again heated by a factor of ten). In what follows, we refer to the fitting procedures that use data augmentation and heated data augmentation as DA and HDA, respectively.

#### From model predictions to forecasts

During each of the CDC weeks DICE was used to fit both the regional and the national most recent incidence data using the combinations of coupling, priors and models described in the previous subsections. The uncoupled procedure (and direct fitting of the National %ILI profile) were used throughout the season with all five priors and with the four models for the force of infection leading to: 5 × 4 = 20 forecasts. For the coupled procedure we fitted with the following combinations of priors and data augmentation: uninformed prior with no data augmentation, uninformed prior with data augmentation and uninformed prior with heated data augmentation (UP, DA and HDA, respectively), and with all four models for the force of infection, leading to: 3 × 4 = 12 forecasts.

This total of 32 model-runs were used to make forecasts of incidence at both the national and regional levels. For each region, we simulate three MCMC chains each with 10^7^ steps and a burn time of 2 × 10^6^ steps. The smallest effective sample size that we report for any parameter was greater than 100. After sampling from the individual posterior densities of each region, we calculated our national forecast as the weighted sum of the regional profiles with the weights given by the relative populations of the regions. The national curve was also fitted directly (without any regional information) using all the models and priors, but these direct results were only used at the end of the season when estimating the performance of each of our procedures.

Early in the season we were experimenting with the coupled procedure and we began to use it as described in the manuscript with the DA and HDA priors only on EW 50 and with the UP prior only on EW 9. Hence, some of the coupled results reported in this section were not available in real-time and were generated at the end of the season (but using only the %ILI data that was available in real-time at each forecast week.)

Each week a single forecast was selected from these results for each of the ten HHS regions and the national. At the regional level we selected a single forecast from one of the (32) uncoupled or coupled procedures enumerated in the previous paragraph. We first reviewed the results for each region and made a selection which took into account: (1) the historic profile for the region, (2) the quality of the fit (both mean and width), (3) the extent of data adjustments in the past weeks, (4) the lab strain % positive data, and when relevant (5) the impact of upcoming school vacations (particularly the winter break). For the national level we selected either one of the 12 coupled results or the aggregated result obtained as the weighted average of our ten regional selections (with the weights given by the relative population of each region). The selection was made by a single group member (and author here) and then reviewed and modified as needed by three group members (also authors here). In most cases (> 90%) there was an agreement on the initial selection. When there was a disagreement, or we were not sure what is the best option, we discussed the selection. All the model results for all seven seasons used here and subsequent seasons are displayed on our website at: http://www.predsci.com/usa-flu-all.

When we evaluate the accuracy of the forecasts we use: (1) The mean absolute error (MAE) of the forecasts (calculated using all 28 weeks of the challenge), (2) The MAE relative to the MAE of a historic NULL model, and (3) The CDC logarithmic score: given a forecast with a set of probabilities for **p**, with *p*_*i*_ being the probability for an observed outcome *p*_*i*_, the logarithmic score is:

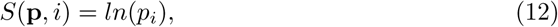

For onset and peak week the score is calculated using the probability assigned to the correct bin plus those of the preceding and proceeding bins (the bin size is one epidemic week). For peak intensity and the 1 − 4 week forward forecast, the score is calculated using the probability assigned to the correct bin plus those of the five preceding and proceeding bins (the bin size is 0.1%).

## Results

### Models selected for forecasts

We selected different FOI variants during different weeks. At the regional level, although we selected the most flexible humidity and school vacation assumptions (HV) more often (47.9% 134/280) than the alternatives (Fig 1), we did select humidity-only (H, 47/280 16.8%), school-vacations-only (V, 62/280 22.1%) and fixed transmissibility (F, 37/280 13.2%) models on a number of occasions. For the national model, we only used the aggregated forecast of the regional models on 9/28 (32.1%) occasions. (The CDC challenge lasts 28 weeks and there are 10 HHS regions, hence the 28 and 280 in the denominators.)

**Fig 1.**
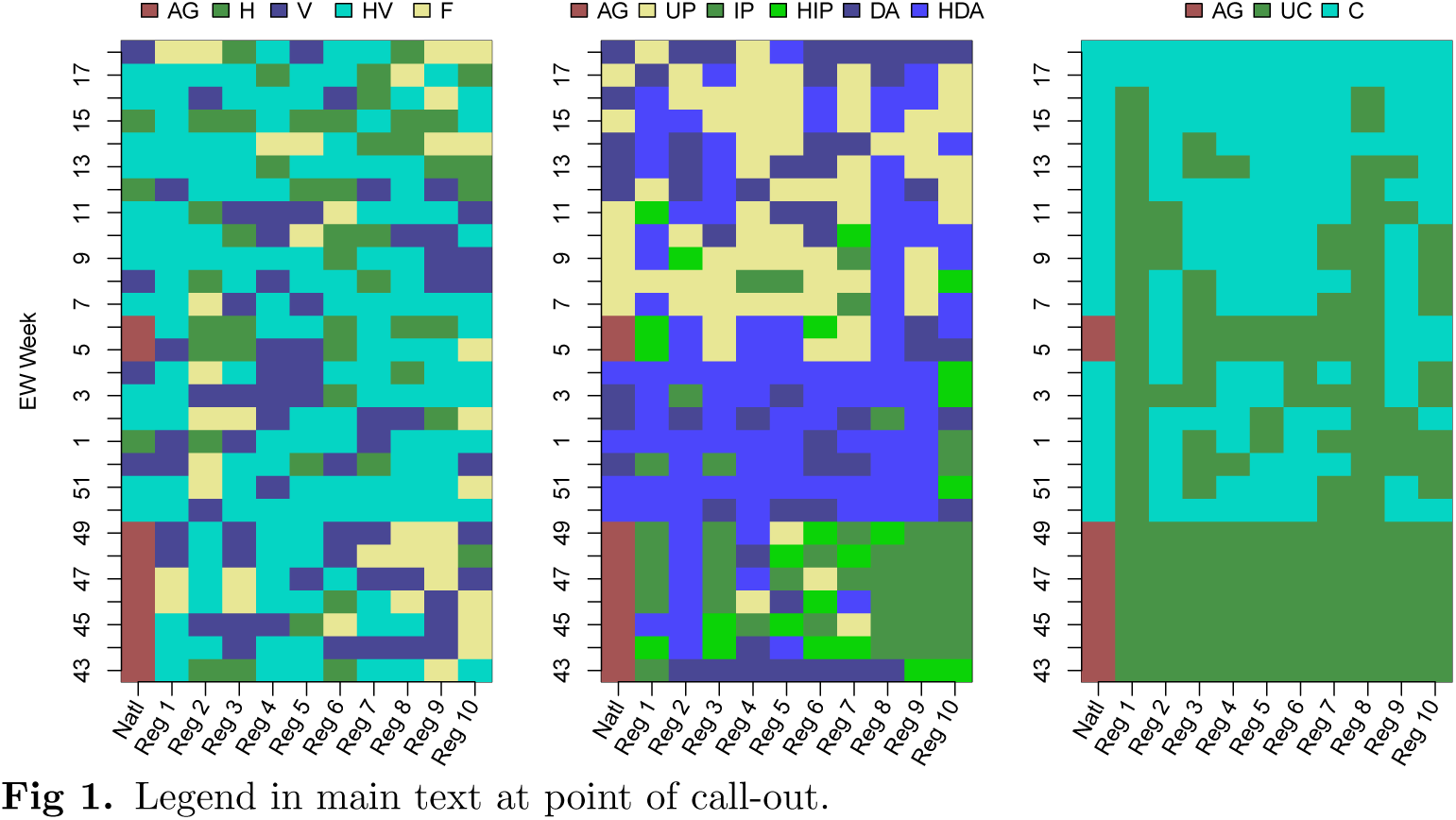
**Forecast model choices for force of infection, use of priors, coupling, and data augmentation; for national and regional estimates by week** Left panel: the selected model for the force of infection: H - specific humidity only, V - school vacation only, HV - both specific humidity and school vacation and F - fixed value for the force of infection. Middle panel: selected model for the prior distribution. UP - uninformed prior, IP - informed prior, HIP - heated informed prior, DA - data augmentation, and HDA - heated data augmentation. Right panel: selected spatial coupling model: UC - uncoupled, C - coupled. For all three panels, AG indicates that the national forecast is aggregated from the regional choices (population-weighted average of the (un)coupled fits to each of the ten HHS regions).

For assumptions about the inferential prior, before the epidemic peaked (EW 06 for the nation) we rarely chose an uninformed prior (UP). Most often we chose the heated data augmentation option (HDA). Early in the season (EWs 43-49), when our options did not include the coupled procedure, we chose the informed prior (IP) or heated informed prior (HIP) most often. Once the season had peaked the uninformed prior was selected often both for the national profile and individual regions, we continued to select the data augmentation (DA) and HDA options for the nation well after the season has peaked (EWs 12-14, 16 and 18).

For assumptions about coupling, once the coupled procedure was available, it was often selected for both the national profile and most regions (1, right panel), with the exception of regions 1 and 8. We found that the coupled procedure used regions 8 (and to a lesser extent 1) as a way to reduce the error to the national fit, at the cost of producing poor fits to these regions, hence their coupled results were rarely selected for submission. The aggregate option for the national selection was only selected at EWs 5 and 6, the weeks prior to the peak and the peak week itself. For these two weeks our errors for both season targets and 1 − 4 week forecasts were large (see Fig 3 below).

### Accuracy of forecasts

At both the national and regional levels, the accuracy of the weekly %-ILI forecasts decreased as the lead time increased. The %ILI 1-4 week forecast and observed data for the national data and the three largest (by population) HHS regions (Fig S1): 4, 5, and 9 is shown in Fig 2. Early in the season, up to and including EW01, the national curve is nearly identical to the historic national curve, whereas our mechanistic forecasts consistently overestimated incidence. After EW01, our national predictions improve significantly, while the historic curve no longer follows the 2016-17 national profile. Similarly, for the three largest HHS regions, the historic curve is similar to the 2016-17 profile until EW01, at which point they start to deviate and our forecasts become more accurate.

**Fig 2.**
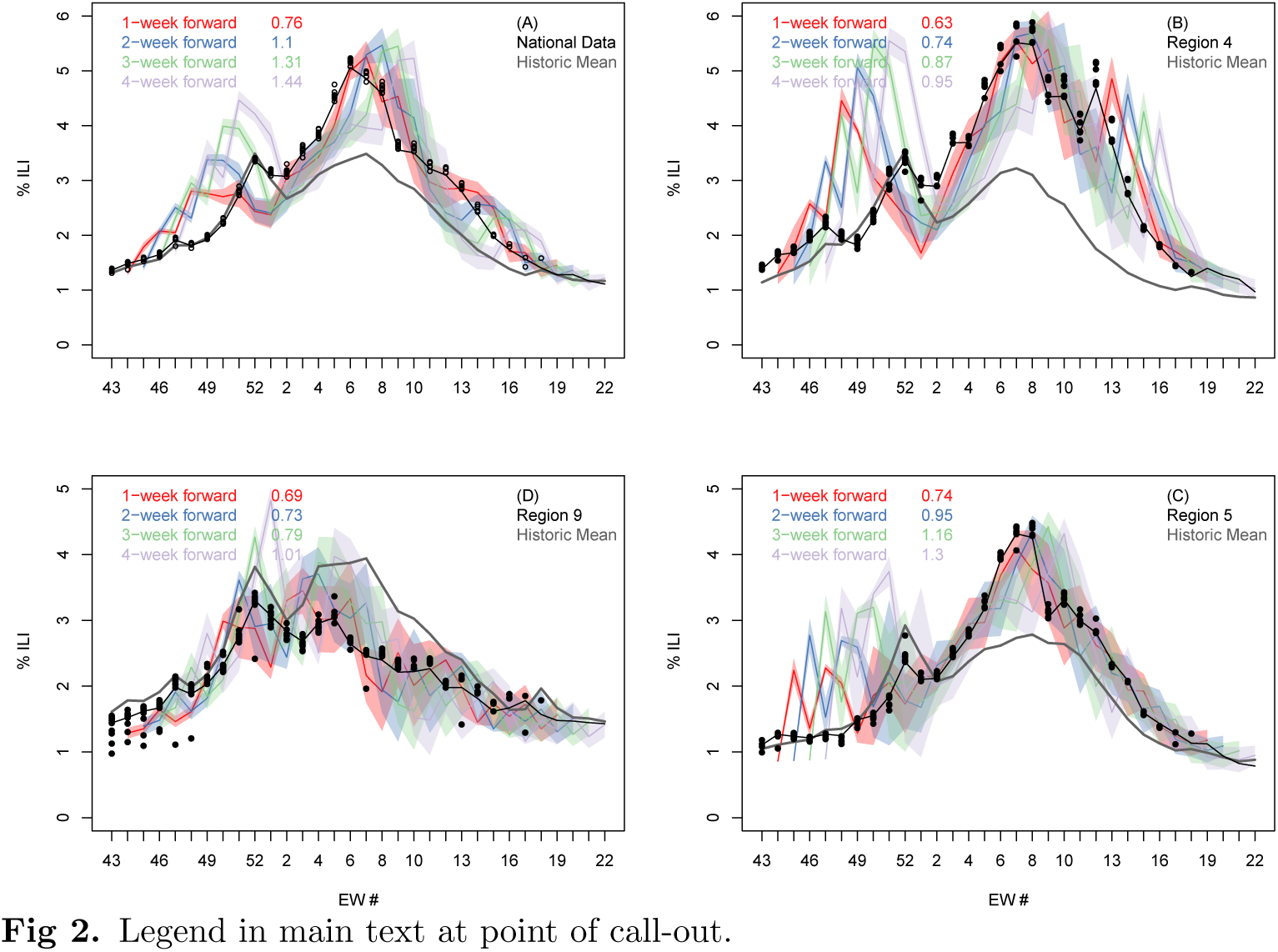
Comparison of submitted forecast and final CDC-reported % of clinic visits that were for ILI for the continental USA and three selected regions. Final season CDC reported (black line), reported during the season (black circles) and predicted %ILI (colored bands) as a function of epidemic week at the national (**A**) and three of the ten regional levels (**B** to **D**). In each panel, the four colored shaded bands denote our one (red), two (blue), three (green) and four (light purple) week-ahead predictions made at each week during the challenge. The width of the colored bands represent our 5-95% prediction intervals as submitted for the forecast. The gray line denotes the historic average, and the numbers in the legend show the error of the DICE forecast relative to the error of the historic NULL model, averaged over all 28 weeks of the challenge. For similar plots for the other seven HHS regions see Fig S2.

Averaged over the entire season, our selected national forecast does better than the historic NULL model only for the 1-week prediction window (Fig 3). However, for the largest HHS region (Region 4) we perform better for all four prediction time horizons, for region 9 (bottom left panel) for the first three, and for region 5 (bottom right panel) for the first two.

**Fig 3.**
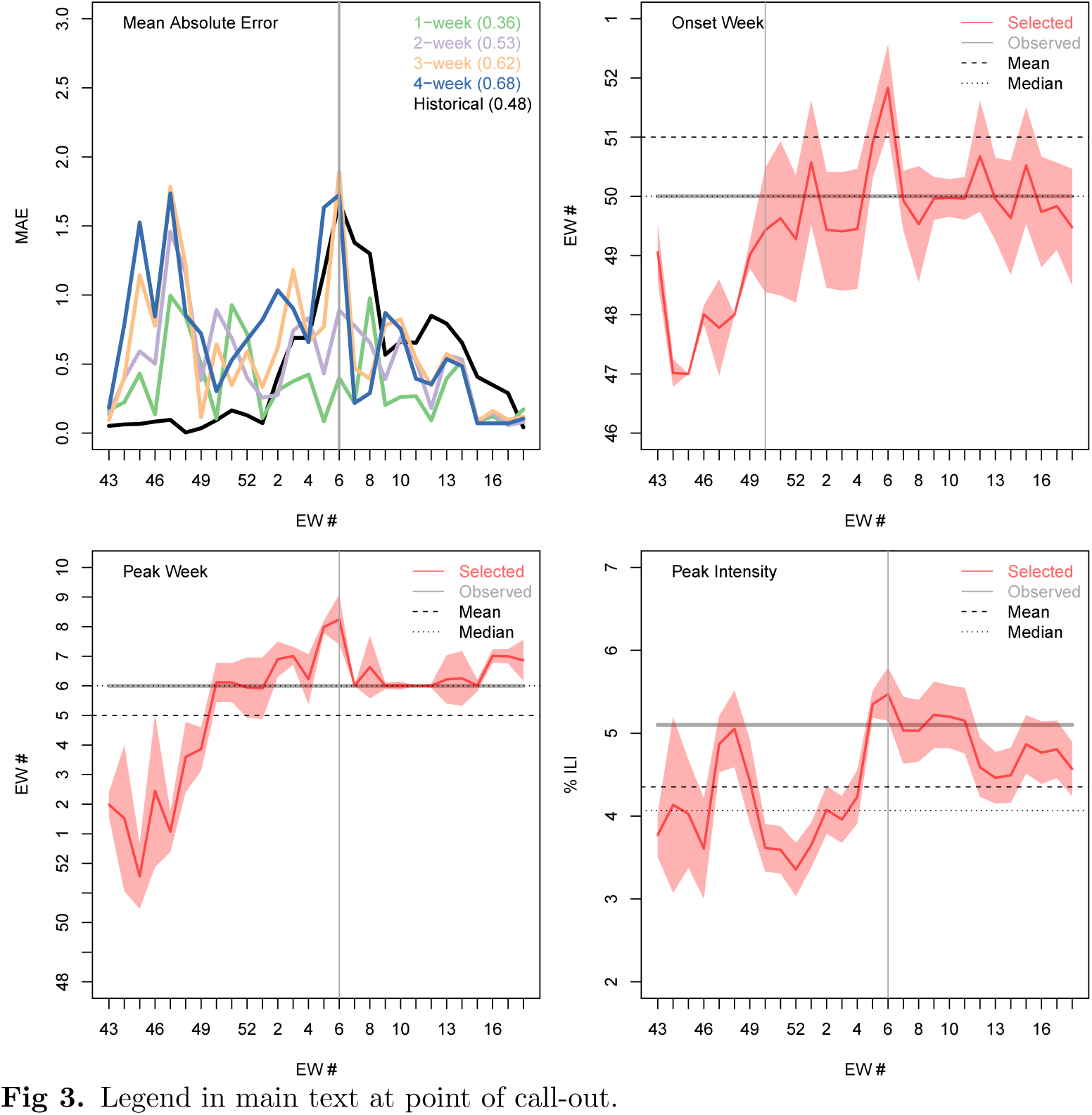
Fig 3. Accuracy of submitted forecasts for timing of onset, timing of peak and intensity of peak. **A** Weekly mean absolute error for the n-week ahead forecast (1 week, green; 2 weeks, purple; 3 weeks, orange; and 4 weeks, blue) and the historic NULL model (black). The average mean absolute error of each forecast horizon, and of the historic prediction, is shown in the legend. Forecast and actual onset week (**B**), peak week (**C**) and peak intensity (**D**) during each week of the CDC challenge: selected forecast (red line); 95% prediction interval (red shading); observed value (gray horizontal line); historic mean (horizontal dashed line); and historic median (horizontal dotted line). The vertical gray line shows the actual 2016-17 peak week (A, C and D) and the actual 2016-17 onset time (B).

Although our forecasts gave potentially useful information over and above the NULL model for the timing of the peak week (Fig 3) and for the amplitude of peak intensity, the peak week of EW06 was the same as the historical mean. Between EW50 (eight weeks before the season peaks) and EW04 (two weeks before the season peak) our forecast correctly predicted to within ±1 week of the observed peak week (EW06). One week before the season peaks, and at the peak week (EW05 and EW06), our model forecast has an error of two weeks.

Forecasts based on the mechanistic model performed better than the historic NULL model for the peak intensity (Fig 3 A/B/C/D). Two weeks before the peak week (and three weeks early in the season) we started predicting the correct peak intensity of 5.1% (to within ±0.5%). The mean and median historic values are significantly lower (4.4% and 4.1% respectively) and outside the ±0.5% range. Our apparent forecast performance for intensity appears to drop off at the end of the season. However, this is an artifact of the forecasting work flow. Once the peak had clearly passed, the final model was selected for reasons other than the peak intensity and the already-observed peak intensity was submitted.

Selected forecasts based on the mechanistic model did not accurately predict onset. Both the mean of onset (EW51) and median (EW50) historic values were within a week of the observed 2016-17 onset week (EW 50). However, our model was unable to properly predict the onset until it happened. As with peak values, once onset had been observed in the data, we used the observed value in our formal submission, which was not reflected in onset values from the chosen model.

Similar models were correlated with each other in their forecasts of 1- to 4-week ahead ILI, but with decreasing strength as forecasts were made for longer time horizons. Fig S5 in the SI shows the Pearson correlation between the 32 models calculated for the 1-, 2-, 3-, and 4- week forward forecast using all 28 weeks of the challenge. The high correlation shows how closely related to each other many of the models are. But this Fig also shows that this correlation decreases when the forecast horizon increases and that there is a spread in the predictions (manifested by negative values of the Pearson correlation).

### Retrospective analyses of model forecasting ability

Once the challenge was over, we examined retrospectively the performance of all mechanistic model variants over the course of all seasons in the historical database and separately for the 2016-17 season. To assess the quality of all the near-term forecasts (1 − 4 weeks) from the different models and assumptions about priors, we show in Fig 4 their weekly CDC score (see Methods), for the 4 different forecast lead times (1-, 2-, 3-and 4- weeks ahead), and for the prediction of the National %ILI intensity. The models are arranged based on their CDC score (averaged across all weeks, numbers on the right y-axis) from best (top) to worse (bottom). Coupled models were more accurate than non-coupled models for the 2016-17 season and for historical seasons for all 4 lead times. The model labeled ‘ensemble’ is the average of the 32 model variants. For the 1-, 2-, and 3-weeks ahead forecasts the subjectively selected model does substantially better than the ensemble model and for the 4-weeks ahead forecast they score practically the same.

**Fig 4.**
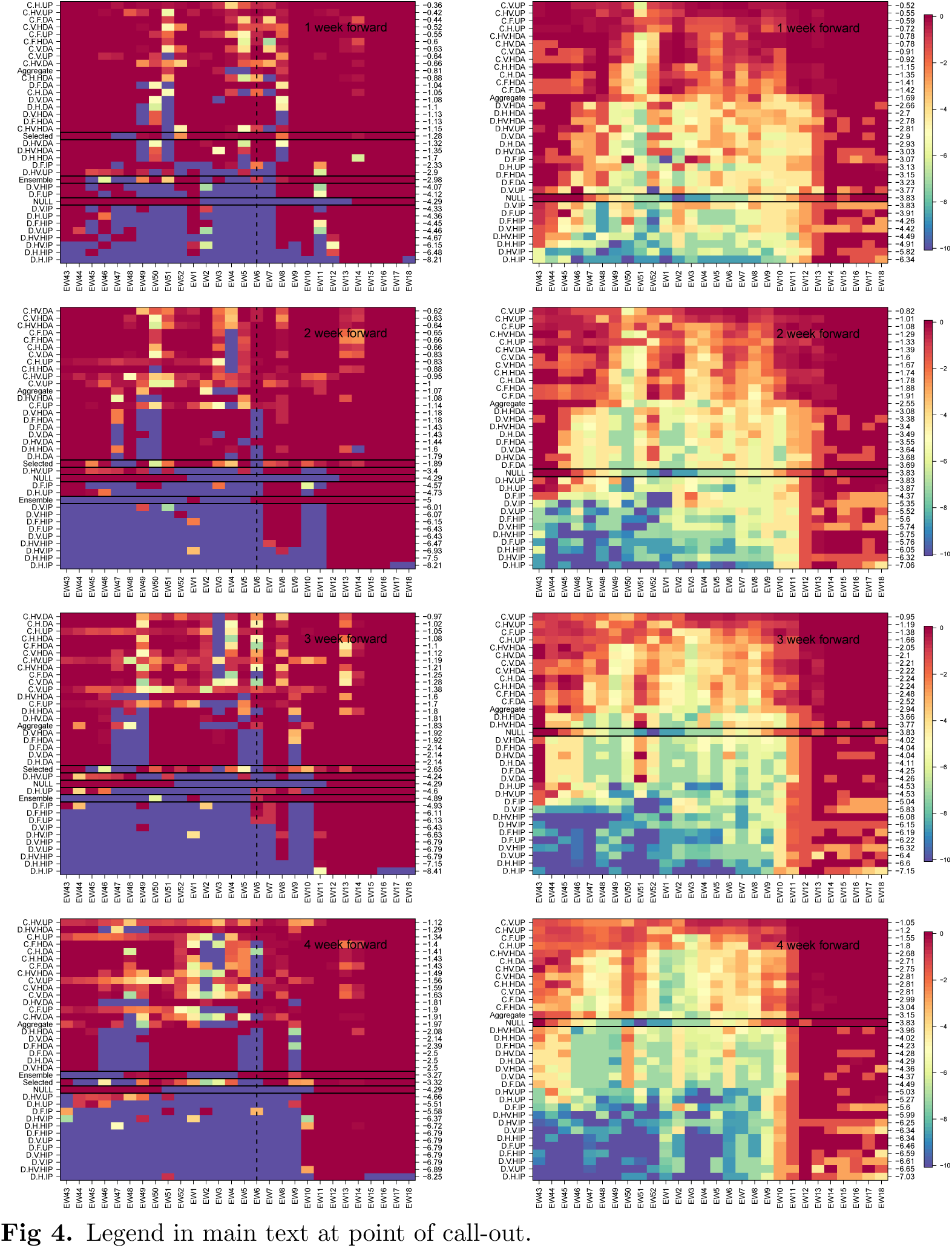
Weekly CDC score for the 1 to 4 week forward national %ILI forecast. Left column: 2016-17 season. Right column: averages over seven flu seasons, 2010-16. In the method/model labels (y-axis): D-direct, C-coupled, and aggregate, H, V, HV and F denote the four models for the force of infection: Humidity only, vacation only, both and fixed. The prior models are: uniform prior (UP), informed prior (IP), heated informed prior (HIP), data augmentation (DA), and heated data augmentation (HDA). The models are ordered by their performance in the season (measured by the mean value of the weekly CDC score) from best (top) to worse (bottom). The mean seasonal score of each model is shown on the right y-axis. Selected (only available for the 2016-17 season), is what we selected each week, the NULL model is the historic average, and ensemble is the average of the 32 model results. A black rectangle is used to highlight these three models. (For the seven season average, right column, we only have the NULL which is highlighted with the black rectangle.) The vertical dashed black line, in the left panels, denotes the national season peak week for the 2016-17 season. For weeks 1-3 the selected model did significantly better than the ensemble and for the 4 weeks forward forecast they performed the same.

The performance of mechanistic models was comparable to that of the historical average NULL model at the beginning and end of the season. However, in the middle of the season, when there is greater variation in the historical data, the performance of the best mechanistic model variants was substantially better than that of the historical average model.

For predicting ILI incidence for the 2016-17 season, which followed similar trend to the historical average, coupled models that used data augmentation were more accurate than coupled models that did not use data augmentation. However, on average for historical seasons, coupled models that did not use augmented data were more accurate than those that did. Also, on average for historical seasons, coupled models that included humidity were more accurate than those that did not (see dark banding in upper portion of charts on the right hand side of Fig 4).

We examined the performance of the different model variants for individual regions for the near-term forecasting of %-ILI (Figs S3 and S4). Again, the coupled models with uninformative prior outperformed other model variants. Although for some regions the improvement in forecast score for the uninformative prior variants over other coupled variants was less pronounced (Regions 1 and 7), these models never appeared to be inferior to the other variants.

In a similar way, we examined the forecast accuracy of different mechanistic model variants in forecasting season-level targets: onset, peak time and peak intensity for the 2016-17 season and on average across all seasons (Fig 5). Here too the models are arranged based on their overall performance from best to worse (top to bottom). Again, for all three targets during 2016-17, the coupled uninformative model variant was at least as good as other coupled options and better than the non-coupled variants. For all three season targets, the selected model performed better (and in the case of peak week significantly better) than the ensemble model. We note that in the latter part of the season, after the single observed onset and peak had passed, results from a single season do not contain much information about model performance. However, the performance of the coupled uninformative prior model was on average better than other model variants across the historical data and different epidemic weeks for all three targets, other than one exception. From EW01 onwards for peak intensity, uncoupled heated augmented prior variants performed better than did coupled uninformative prior model variants.

**Fig 5.**
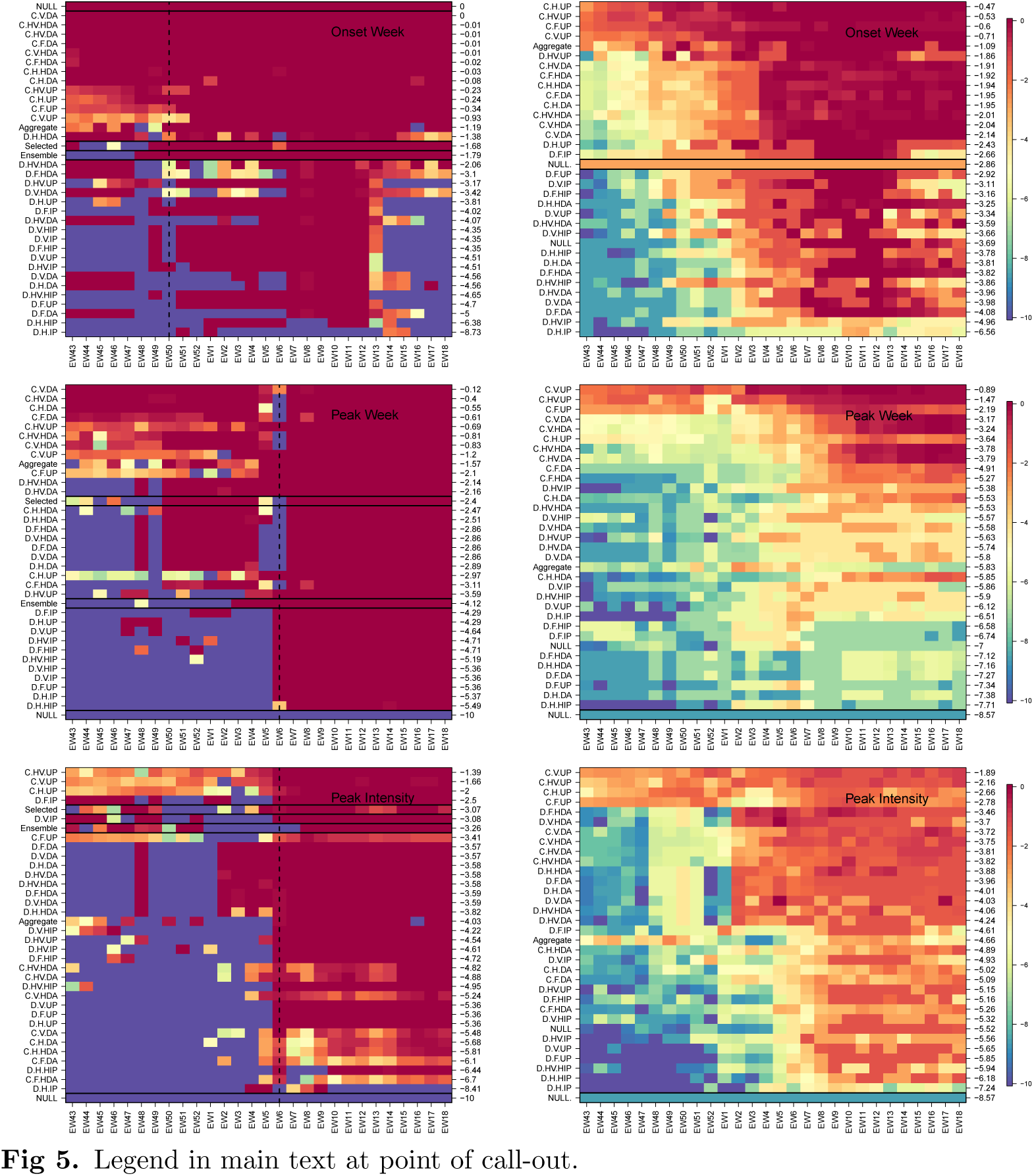
Weekly CDC score for season targets: timing of onset, timing of peak and intensity of peak. As in Fig 4 but for the seasonal targets: onset week (top), peak week (middle) and peak intensity (bottom). Left column: 2016-17 season, right column: averaged over seven seasons, 2010-16. Models are arranged from best to worse (top to bottom) and the mean season score for each model is shown on the right y-axis. The ensemble model is the average of the 32 model variants. In the left column panels, the selected, ensemble and NULL models are denoted with a black rectangle. The latter is also highlighted in the right column. For the 2016-17 season, the coupled procedure with data augmentation correctly predicts onset week and peak week from the start of the CDC challenge. Peak intensity is predicted best with a coupled uninformed prior. When averaged over seven seasons (right column) the coupled uninformed prior does best for all three season targets.

We were able to compare our performance over the course of the season to the performance of the other teams using the public website that supports the challenge (www.cdc.gov/flusight). Averaged across all weeks of forecast and all forecast targets, we were ranked 13 out of 29 teams. For 1-, 2-, 3- and 4-weeks ahead forecasts we were ranked 6, 11, 9 and 16 respectively; again, out of 29 teams. We were ranked 14 for the timing of onset, 5 for the timing of the peak and 14 for intensity of the peak. Probing beyond the overall rankings, our performance was similar to the other better-performing teams in the challenge. Also, our performance improved substantially as measured by both in absolute terms and relative to other teams across the season (Figs S6 and S7).

## Discussion

In this study, we have described our participation in a prospective forecasting challenge. Although we drew on results from a large set of mechanistic models, our single forecast for each metric was made after choosing between available model results for that metric in that week and was therefore somewhat subjective. We performed poorly at the start of the competition when our mechanistic models consistently over-estimated incidence. However, during the middle phase of the season, our models produced less biased estimates and consistently outperformed non-mechanistic models based on the average of historical data. A robust testing of model variants using historical data suggests that spatially coupled models are systematically better than historical NULL models during the middle of the season and are not significantly worse even at the start of the season. We evaluated a simple ensemble and showed that the subjective model choice was better. However, the ranking of individual models suggests that an ensemble of coupled models may outperform our subjective choice. We are considering exactly this experiment for the upcoming season.

This study is slightly different from some prior studies of influenza forecasting [30] in that it describes and assesses a subjective choice between multiple mechanistic models as the basis of a prospective forecast, rather than describing the performance of a single model or single ensemble of models used for an entirely objective forecast. Although this could be viewed as a limitation of our work, because individual subjective decisions cannot be reproduced, we suggest that the explicit description of a partially subjective process is a strength. In weather forecasting, there is a long history of evaluating the accuracy of entirely objective forecasts versus partially subjective forecasts [31,32]. Broadly, for each different forecast target and each forecast lead-time, there has been a gradual progression over time such that objective forecasts become more accurate than subjective forecasts. We note also that although we describe the subjective process as it was conducted, we also provide a thorough retrospective assessment of the predictive performance of each model variant.

We may refine our ensemble approach for future iterations of the competition. It seems clear that the coupled models produce more accurate forecasts than the uncoupled models for most targets, so we would consider an ensemble only of the coupled model variants. We will also consider weighted ensembles of models and attempt to find optimal weights by studying all prior years. Also, data were often updated after being reported and we did not include an explicit reporting model in our inferential framework (also sometimes referred to as a backfill model). Rather, we used knowledge of past adjustments to data during our discussions and eventual subjective choice of models. We aim to include a formal reporting model in future versions of our framework.

By reflecting on our choices and their performance, we can evaluate the importance of a number of different model features. Our coupled model variants performed much better than uncoupled variants consistently across the 2016-17 season, for different targets and when evaluated using the historical data. This prospective study supports recent retrospective results suggesting that influenza forecasts can be more accurate if they explicitly represent spatial structure [33,34]. Given that the model structure we used to represent space was relatively coarse [24], further work is warranted to test how forecast accuracy at finer spatial scales can be improved by models that include iteratively finer spatial resolution.

In submitting forecasts based on uninformed mechanistic priors using an uncoupled model at the start of the season, we failed to learn lessons that have been present in the influenza forecasting literature for some time [30]. Historical variance is low during the start of the season and the growth pattern is not exponential. Therefore, it would be reasonable to forecast early exponential growth only in the most exceptional of circumstance, such as during the early weeks of a pandemic. Model solutions that are anchored to the historical average in some way, such by the use of augmented data for not-yet-seen time points, are likely to perform better. Also, forecasting competitions may want to weight performance differentially across time, with greater weight given to forecasts during periods where there is a higher variance in incidence.

Models that included humidity forcing performed better on average in our analysis of all historical data than equivalent models that did not include those terms, especially for the forecasting of ILI 1- to 4-weeks ahead [35]. However, we did not see similar support for the inclusion of school vacation terms improving accuracy, which has been suggested in a retrospective forecasting study at smaller spatial scales (by this group) [36]. The lack of support for school vacations in the present study could indicate that the prior work was under-powered or that averaging of school vacation effects across large spatial scales – both in the data and the model – degrades its contribution to forecast accuracy. Also, we chose to present our accuracy of predicting peak height and timing relative to actual epidemiological week so that it was consistent with our presentation of accuracy of other forecast targets. However, it may be more appropriate in some circumstances to present accuracy of targets associated with the peak relative to the eventual peak [11].

We found the experience of participating in a prospective forecasting challenge to be different to that of a retrospective modeling study. The feedback in model accuracy was much faster and the need for statistically robust measures of model likelihood or parsimony less obvious. We encourage the use of forecasting challenges for other infectious disease systems as a focus for better understanding of underlying dynamics in addition to the generation of actionable public health information.

## Acknowledgments

Disclaimer: The findings and conclusions in this report are those of the author(s) and do not necessarily represent the views of the Department of Health and Human Services or its components, the US Department of Defense, local country Ministries of Health, Agriculture, or Defense, or other contributing network partner organizations. Mention of any commercial product does not imply DoD endorsement or recommendation for or against the use of any such product. No infringement on the rights of the holders of the registered trademarks is intended. No funding bodies had any role in study design, data collection and analysis, decision to publish, or preparation of the manuscript.

## Supporting information

**S1 Text. Details of the parametric dependence of the force of infection on specific humidity**

**S2 Text. Details of the parametric dependence of the force of infection on school vacation schedule**

**S1 Fig.**
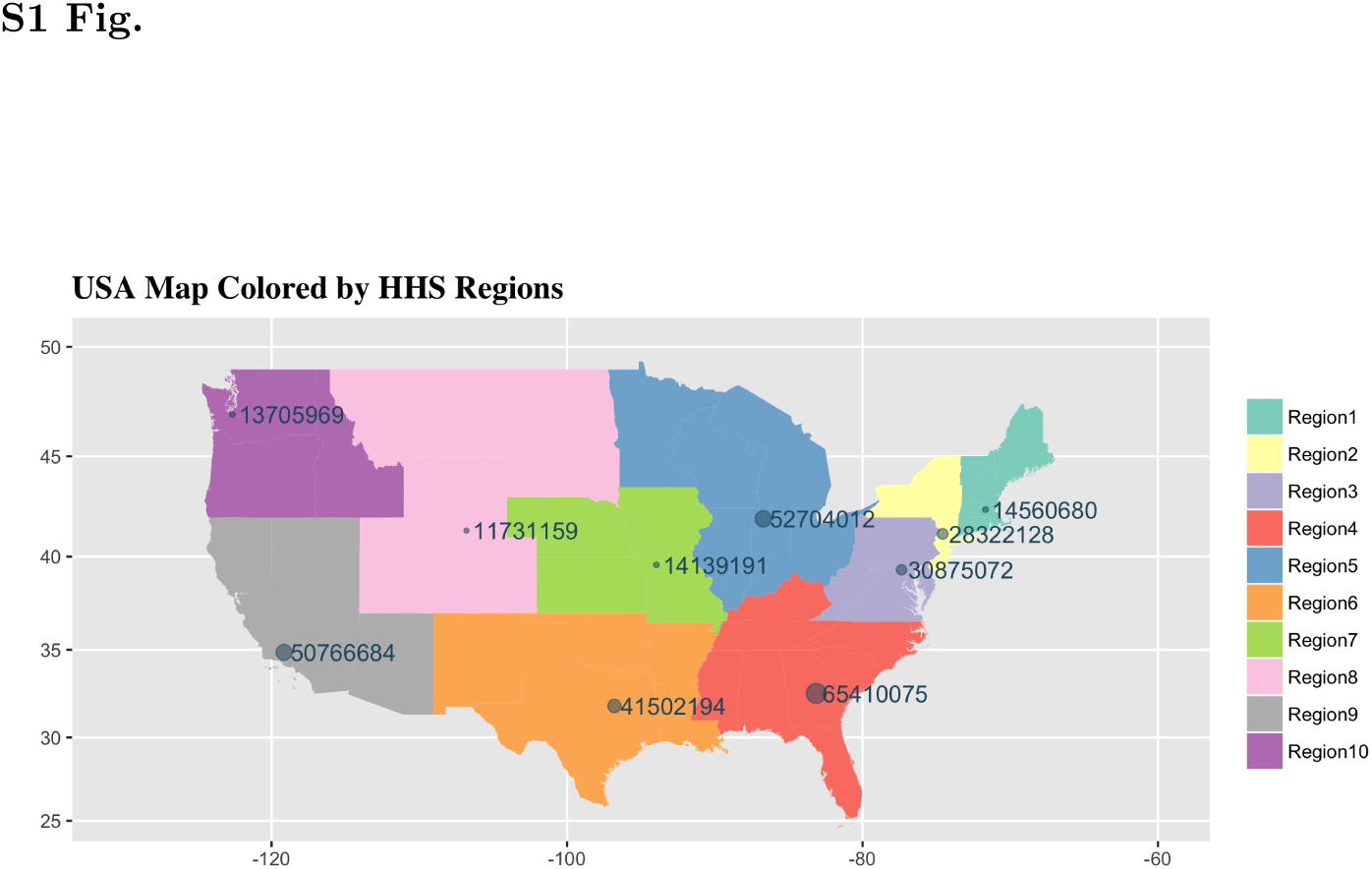
A map of the contiguous US colored by the ten HHS regions

The green circle in each HHS region denotes the population density weighted location of the centroid of the region, and the radius of each circle is proportional to the weight of the region which is determined by its relative population. The population of each region is denoted on the map next to the centroid.

**S2 Fig.**
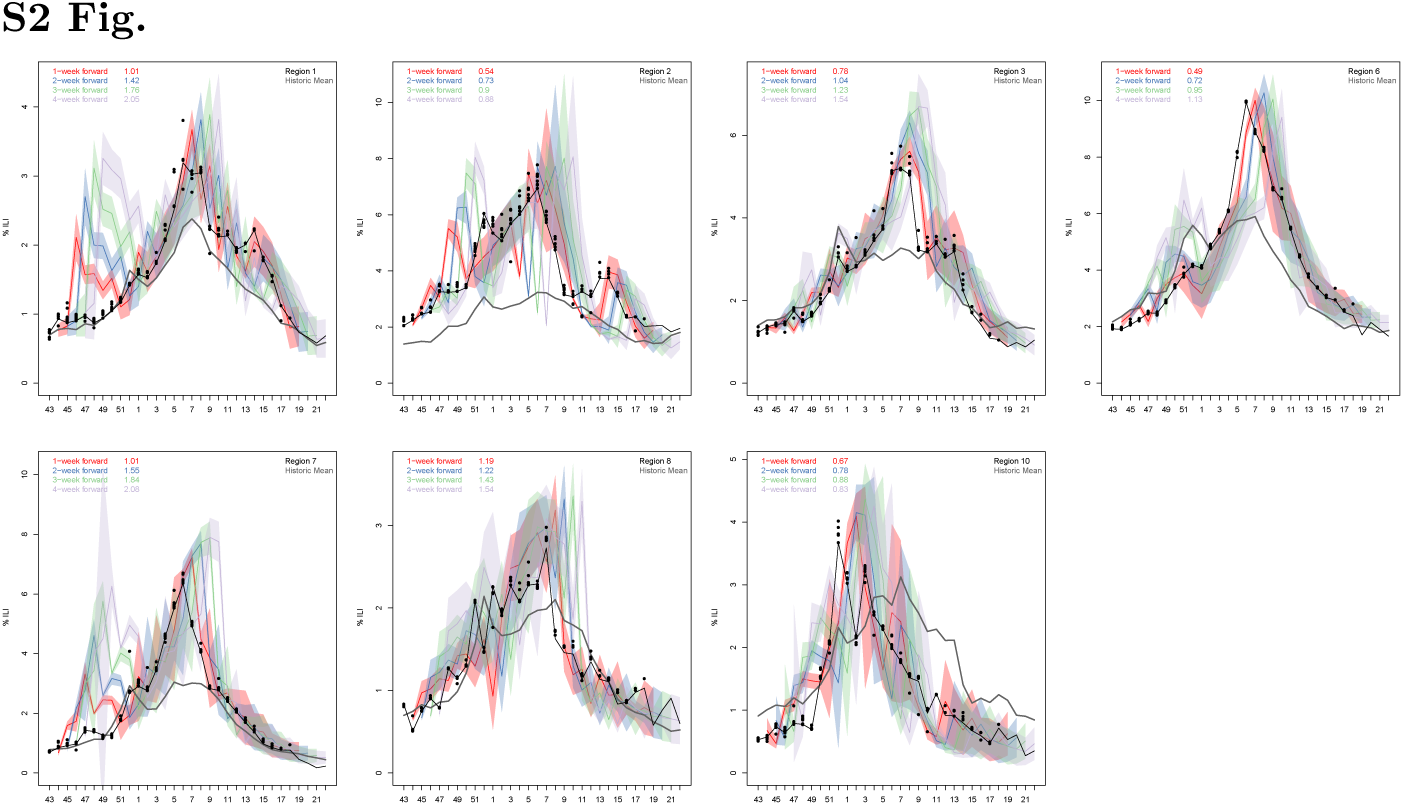
Predicted %ILI as a function of epidemic week for seven of the ten HHS regions.

Final season CDC reported (black line), reported during the season (black circles) and predicted %ILI (colored bands) as a function of epidemic week for seven of the ten HHS regions not shown in the main text panel. In each panel the four colored shaded bands denote our *n*-week forward prediction (*n* = 1, 2, 3, 4), and the gray line denotes the historic average. The average relative error (measured with respect to the error of the historic NULL model) is indicated for each of the four prediction horizons in the legend. See main text and Fig. 2 for more details.

**S3 Fig. Zoom-able separate file of weekly CDC score for** 1 − 4 **week forward regional forecast averaged over the 2010-16 seasons.**

As in the right columns of Fig 4 but for the ten HHS regions. Each columns denotes an HHS region and going down a columns we move from 1 to 2, 3 and 4 weeks forecasts. In the method/model labels (y-axis): UN-uncoupled and C-coupled. H, V, HV and F denote the four models for the force of infection: Humidity only, vacation only, both and fixed. The prior models are: uniform prior (UP), informed prior (IP), heated informed prior (HIP), data augmentation (DA), and heated data augmentation (HDA). Selected, is what we selected each week, the NULL model is the historic average and the ensemble is the average of the 32 model variants. Models are arranged based on their overall performance during the entire season (numbers on the right y-axis) from best (top) to worse (bottom). For all ten regions, and in agreement with the results for the nation, the coupled procedure performs better than the uncoupled. For nearly all regions and forecasts lead times the selected option does better than the ensemble model. To view this Figure better please use the ‘zoom’ tool when opening it with a PDF viewer.

**S4 Fig. Zoom-able separate file of weekly CDC score for the seasonal targets: onset week, peak week and peak intensity for the ten HHS regions averaged over the 2010-16 seasons.**

Top, middle and bottom rows are onset week, peak week and peak intensity. The NULL result is calculated using the historic mean regional profile. The ensemble mode is the average of the 32 model variants. For all three targets, and all ten regions, the coupled method does better than the uncoupled with the details of the prior and force of infection models depending on the region. For nearly all targets and regions the selected model does better than the ensemble model. To view this Figure better please use the ‘zoom’ tool when opening it with a PDF viewer.

**S5 Fig.**
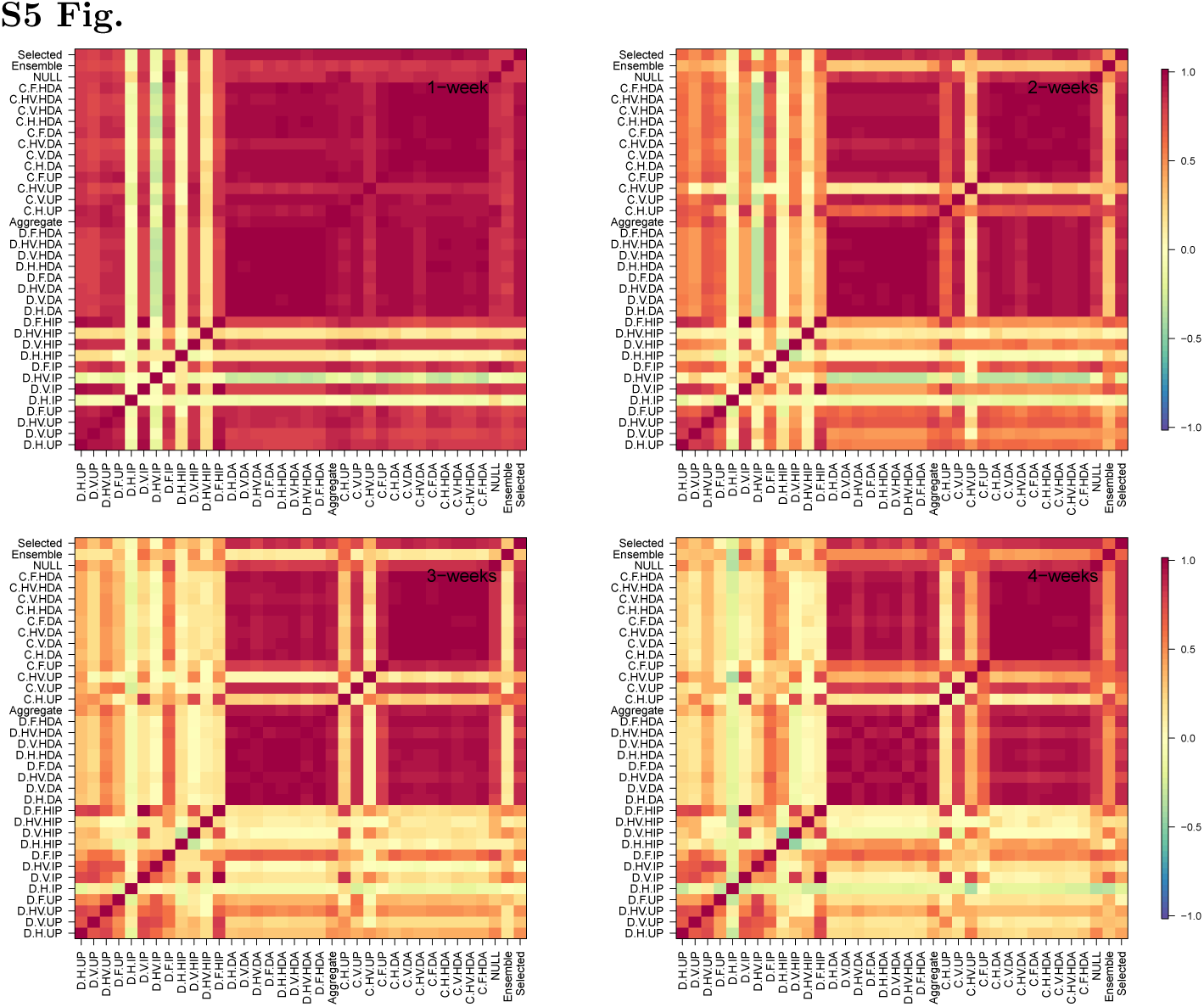
**Correlation between the** 1 − 4 **week forward nation forecast**

The Pearson correlation between the 1-, 2-, 3- and 4-week forward nation forecast calculated using all models and all 28 weeks of the 2016-17 CDC challenge. For the 1 − 4 weeks forward forecasts the correlation within coupled models and uncoupled models is greater than between uncoupled and coupled. As the prediction horizon increases the correlation between models decreases significantly, but it remains high for the coupled models.

**S6 Fig.**
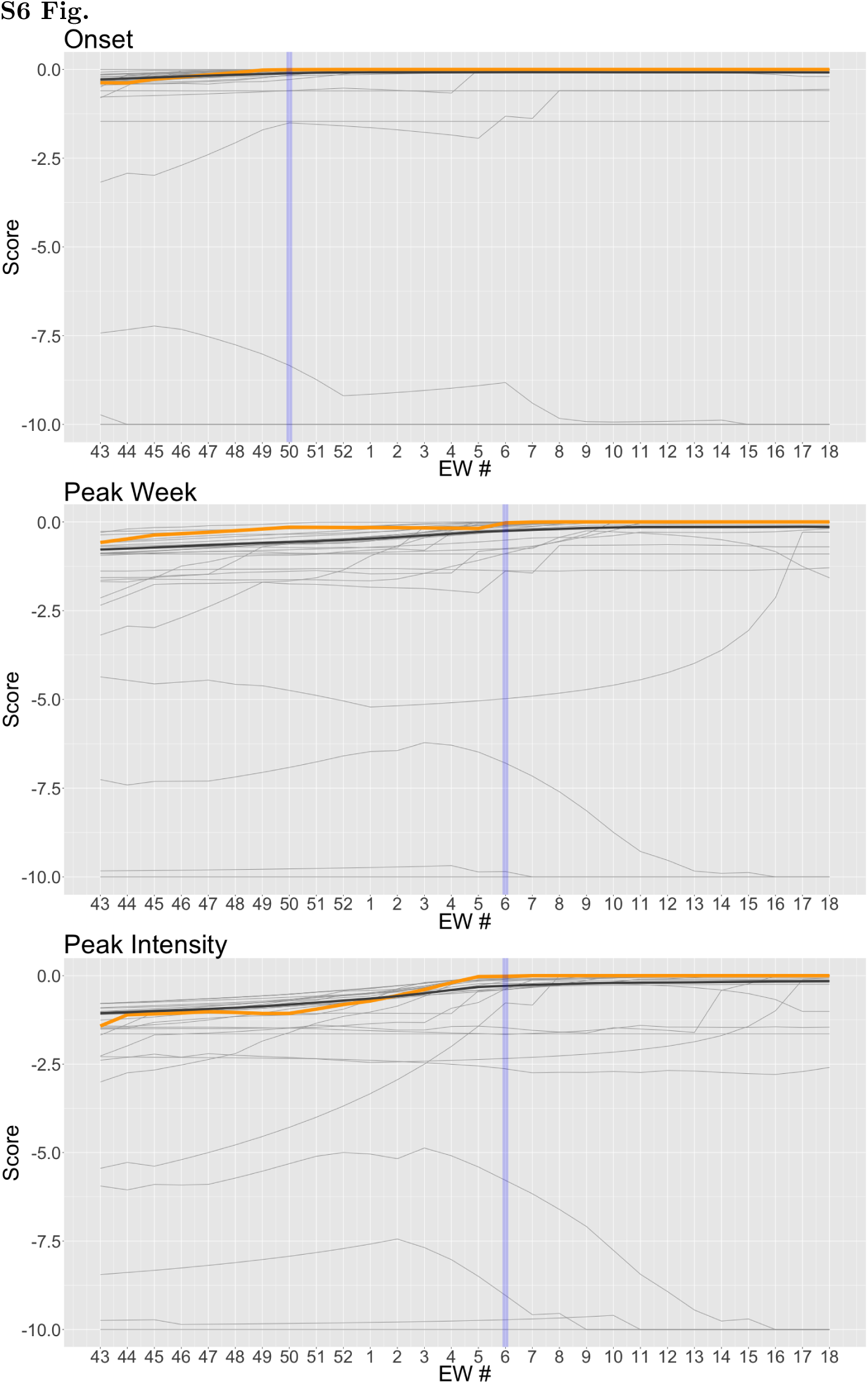
Comparison of 2016-17 weekly reverse rolling scores for all submissions: Nation season targets.

Top, middle and bottom rows are onset week, peak week and peak intensity. Each light gray line denotes one of the 29 submissions to the CDC challenge, the black line in the unweighted average of these submissions and the orange line is that of the DICE model submission. Here, the weekly score is calculated as a reverse rolling average where the first week (EW 43) is the average of all 28 weeks of the challenge, the second week averages the score from week 2 (EW 44) onward etc. This reverse rolling average highlights the gradual improvement of the DICE submission. The vertical blue line marks the season onset week (top panel) and peak week (middle and bottom) panels.

**S7 Fig.**
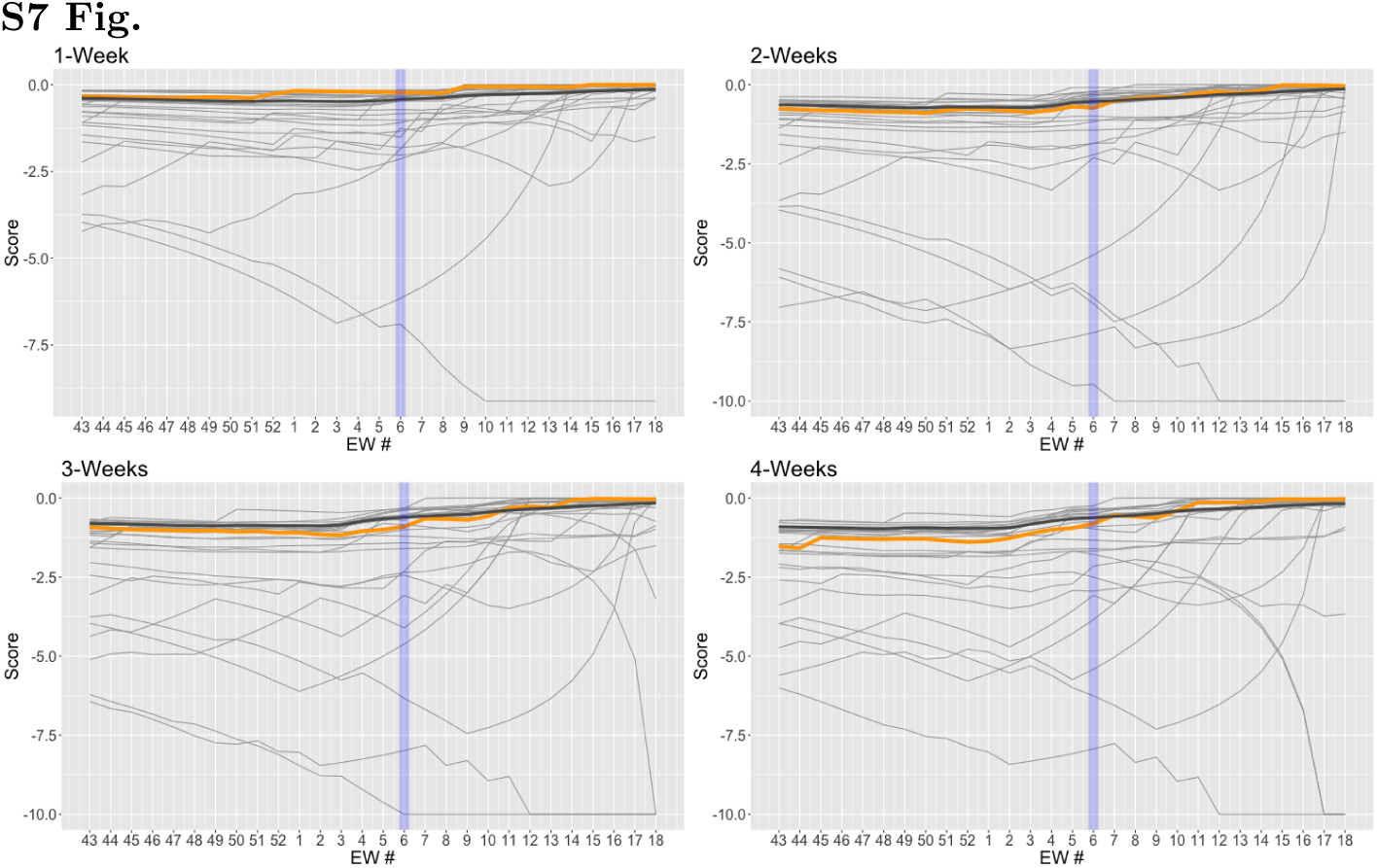
**Compsrison of 2016-17 weekly reverse rolling scores for all submissions:** 1 − 4 **Week forward nation forecast.**

Similar to Fig S6, but for the 1- to 4- week forward forecast.

**S1 Text Details of the parametric dependence of the force of infection on specific humidity**

Eq. 11 allows the transmission rate to depend on time using three terms. The first time dependent term, *F*_1_(*t*), allows for a dependence of the transmission rate on specific-humidity. In temperate regions specific humidity has a seasonal oscillation with a minimum in the winter and a maximum in the summer. We follow Shaman et al. [37] and relate the local SH, *q*_*j*_(*t*), to the reproduction number as:

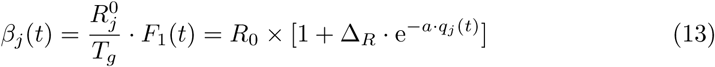

In the above equation, and unlike the work published by others, the values of the parameters *a* and Δ_*R*_ are fitted. The effect of specific humidity can be combined with that of school vacation which is discussed in the following sub-section.

**S2 Text Details of the parametric dependence of the force of infection on school vacation schedule**

The second term in Eq. 11 allows the transmission rate to depend on the weekly school vacation schedule (*p*_*j*_(*t*)) and we implement is as:

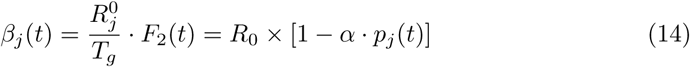

DICE fits the effect of school closure by optimizing the parameter *α*, which is in the range 0 − 1. Larger values of *α* indicate a more significant lowering of *β*_*j*_(*t*) as a result of planned school vacations. Conversely, small values of *α* indicate that the school vacation schedule is not an important factor in determining the ILI profile. The effect of school vacation can be combined with that of specific humidity, i.e. 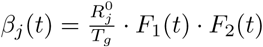

## References

1. WHO Seasonal Influenza; accessed October 9, 2018. http://www.who.int/news-room/fact-sheets/detail/influenza-(seasonal)t.

2. Tsang KW, Ho PL, Ooi GC, Yee WK, Wang T, Chan-Yeung M, et al. A cluster of cases of severe acute respiratory syndrome in Hong Kong. N Engl J Med. 2003;348(20):1977–1985.

3. Centers for Disease Control and Prevention CDC. Swine influenza A (H1N1) infection in two children–Southern California, March-April 2009. MMWR Morb Mortal Wkly Rep. 2009;58(15):400–402.

4. Campos GS, Bandeira AC, Sardi SI. Zika Virus Outbreak, Bahia, Brazil. Emerg Infect Dis. 2015;21(10):1885–1886.

5. Chertien JP, George D, Shaman J, Chitale RA, E MF. Influenza Forecasting in Human Populations: a Scoping Review; 2014. Available from: https://doi.org/10.1371/journal.pone.0094130.

6. Hanratty B, Robinson M. Coping with winter bed crises. New surveillance systems might help. BMJ. 1999;319(7224):1511–1512.

7. Biggerstaff M, Johansson M, Alper D, Brooks LC, Chakraborty P, Farrow DC, et al. Results from the second year of a collaborative effort to forecast influenza seasons in the United States. Epidemics. 2018;24. doi:10.1016/j.epidem.2018.02.003.

8. Brooks LC, Farrow DC, Hyun S, Tibshirani RJ, Rosenfeld R. Nonmechanistic forecasts of seasonal influenza with iterative one-week-ahead distributions. PLOS Computational Biology. 2018;14(6).

9. Ray EL, Krzysztof S, Lauer SA, Johansson MARNG. Infectious disease prediction with kernel conditional density estimation. Statistics in Medicine. 2017;.

10. Ray EL, G RN. Prediction of infectious disease epidemics via weighted density ensembles. PLOS Computational Biology. 2018;14(2).

11. Zhang Q, Perra N, Perrotta D, Tizzoni M, Paolotti D, A V. Forecasting seasonal influenza fusing digital indicators and a mechanistic disease model. In: Proceedings of the 26th International Conference on World Wide Web; 2017. p. 311–319.

12. Tizzoni M, Bajardim P, Poletto C, Balcan D, Goncalves B, Perra N, et al. Real-time numerical forecast of global epidemic spreading: case study of 2009 A/H1N1pdm. BMC Medicine;10(1). doi:10.1186/1741-7015-10-165.

13. Osthus D, Gattiker J, Priedhorsky R, Valle SYD. Dynamic Bayesian Influenza Forecasting in the United States with Hierarchical Discrepancy; 2017.

14. Pei S, J S. Counteracting structural errors in ensemble forecast of influenza outbreaks. Nature Communications. 2017;8(1). doi:10.1038/s41467-017-01033-1.

15. Yang W, Karspeck A, J S. Comparison of Filtering Methods for the Modeling and Retrospective Forecasting of Influenza Epidemics. PLOS Computational Biology. 2014;10(4).

16. Olson DR, Konty KJ, Paladini M, Viboud C, Simonsen L. Reassessing Google Flu Trends data for detection of seasonal and pandemic influenza: a comparative epidemiological study at three geographic scales. PLoS Comput Biol. 2013;9(10):e1003256.

17. Centers for Disease Control and Prevention, CDC: Flu Activity & Surveillance; accessed September 14, 2016. http://gis.cdc.gov/grasp/fluview/fluportaldashboard.html.

18. Schappert S, Burt C. Ambulatory care visits to physician offices, hospital outpatient departments, and emergency departments: United States, 2001-02. Vital and Health Statistics Series 13, Data from the National Health Survey. 2006;(159):1–66.

19. Talwalkar A, Hing E, Palso K. National Ambulatory Medical Care Survey: 2011 Summary Tables;. Available from: http://www.cdc.gov/nchs/ahcd/ahcd_products.htm.

20. Talwalkar A, Hing E, Palso K. National Hospital Ambulatory Medical Care Survey: 2011 Outpatient Department Summary Tables;. Available from: http://www.cdc.gov/nchs/ahcd/ahcd_products.htm.

21. National Aeronautics and Space Administration (NASA): Land Data Assimilation Systems; accessed September 15, 2016. http://ldas.gsfc.nasa.gov/nldas/NLDAS2forcing.php.

22. Xia Y, Mitchell K, Ek M, Sheffield J, Cosgrove B, Wood E, et al. Continental-scale water and energy flux analysis and validation for the North American Land Data Assimilation System project phase 2 (NLDAS-2): 1. Intercomparison and application of model products. J Geophys Res-Atmos. 2012;117(D3).

23. Mitchell KE, Lohmann D, Houser PR, Wood EF, Schaake JC, Robock A, et al. The multi-institution North American Land Data Assimilation System (NLDAS): Utilizing multiple GCIP products and partners in a continental distributed hydrological modeling system. J Geophys Res-Atmos. 2004;109(D7).

24. Mills HL, Riley S. The Spatial Resolution of Epidemic Peaks. PLoS Comput Biol. 2014;10(4):1–9.

25. Riley P, Ben-Nun M, Armenta R, Linker JA, Eick AA, Sanchez JL, et al. Early Characterization of the Severity and Transmissibility of Pandemic Influenza Using Clinical Episode Data from Multiple Populations. PLoS Comput Biol. 2013;9(5):1–15. doi:10.1371/journal.pcbi.1003064.

26. Riley P, Ben-Nun M, Linker JA, Cost AA, Sanchez JL, George D, et al. Early Characterization of the Severity and Transmissibility of Pandemic Influenza Using Clinical Episode Data from Multiple Populations. PLoS Comput Biol. 2015;11(9):1–15. doi:10.1371/journal.pcbi.1004392.

27. Gilks WR, Richardson S, Spiegelhalter DJ. Markov Chain Monte Carlo in Practice. Chapman and Hall/CRC Interdisciplinary Statistics Series. Chapman & Hall; 1996. Available from: http://books.google.com/books?id=TRXrMWY_i2IC.

28. Moss R, Zarebski AF, Dawson P, McCaw JM. Retrospective forecasting of the 2010–2014 Melbourne influenza seasons using multiple surveillance systems. Epidemiology and Infection;145:156–169. doi:10.1017/S0950268816002053.

29. Zarebski AF, Dawson P, McCaw JM, Moss R. Model Selection for Seasonal Influenza Forecasting. Infectious Disease Modelling;2:56–70. doi:https://doi.org/10.1016/j.idm.2016.12.004.

30. Chretien JP, Dylan G, Shaman J, Chitale RA, McKenzie FE. Influenza Forecasting in Human Populations: A Scoping Review. PLoS ONE;9. doi:https://doi.org/10.1371/journal.pone.0094130.

31. Murphy AH, Brown BG. A comparative evaluation of objective and subjective weather forecasts in the united states. J Forecast. 1984;3(4):369–393.

32. Murphy AH, Winkler RL. Probability Forecasting in Meteorology. J Am Stat Assoc. 1984;79(387):489–500.

33. Pei S, Kandula S, Yang W, Shaman J. Forecasting the spatial transmission of influenza in the United States. Proc Natl Acad Sci U S A. 2018;.

34. Yang W, Olson DR, Shaman J. Forecasting Influenza Outbreaks in Boroughs and Neighborhoods of New York City. PLoS Comput Biol. 2016;12(11):e1005201.

35. Axelsen JB, Yaari R, Grenfell BT, Stone L. Multiannual forecasting of seasonal influenza dynamics reveals climatic and evolutionary drivers. Proc Natl Acad Sci U S A. 2014;111(26):9538–9542.

36. Riley P, Ben-Nun M, Turtle JA, Linker J, Bacon DP, Riley S. Identifying factors that may improve mechanistic forecasting models for influenza. bioRxiv. 2017;doi:10.1101/172817.

37. Shaman J, Pitzer VE, Viboud C, Grenfell BT, Lipsitch M. Absolute Humidity and the Seasonal Onset of Influenza in the Continental United States. PLoS Biol. 2010;8(2):1–13.

